# Yeast cells actively tune their membranes to phase separate at temperatures that scale with growth temperatures

**DOI:** 10.1101/2021.09.14.460156

**Authors:** Chantelle L. Leveille, Caitlin E. Cornell, Alexey J. Merz, Sarah L. Keller

**Affiliations:** Department of Chemistry, University of Washington, Seattle WA, USA; Department of Biochemistry, University of Washington, Seattle WA, USA

**Keywords:** phase separation, yeast, vacuole, membrane, raft

## Abstract

Membranes of vacuoles, the lysosomal organelles in yeast, undergo extraordinary changes during the cell’s normal growth cycle. The cycle begins with a stage of rapid cell growth. Then, as glucose becomes scarce, growth slows, and the vacuole membranes phase-separate into micron-scale liquid domains. Recent studies suggest that these domains are important for yeast survival by laterally organizing membrane proteins that play a key role in a central signaling pathway conserved among eukaryotes (TORC1). An outstanding question in the field has been whether yeast stringently regulate the phase transition and how they respond to new physical conditions. Here, we measure transition temperatures – an increase of roughly 15°C returns vacuole membranes to a state that appears uniform across a range of growth temperatures. We find that broad populations of yeast grown at a single temperature regulate the transition to occur over a surprisingly narrow temperature range. Moreover, the transition temperature scales linearly with the growth temperature, demonstrating that the cells physiologically adapt to maintain proximity to the transition. Next, we ask how yeast adjust their membranes to achieve phase separation. Specifically, we test how levels of ergosterol, the main sterol in yeast, induce or eliminate membrane domains. We isolate vacuoles from yeast during their rapid stage of growth, when their membranes do not natively exhibit domains. We find that membrane domains materialize when ergosterol is depleted, contradicting the assumption that increases in ergosterol cause membrane phase separation *in vivo*, and in agreement with prior studies that use artificial and cell-derived membranes.

**SIGNIFICANCE STATEMENT:** Phase separation in membranes creates domains enriched in specific components. To date, the best example of micron-scale phase separation in the membrane of an unperturbed, living cell occurs in a yeast organelle called the vacuole. Recent studies indicate that the phases are functionally important, enabling yeast survival during periods of cellular stress. We have discovered that yeast regulate this phase transition; the temperature at which membrane components mix into a single phase is ~15°C above the growth temperature. To maintain this offset, yeast may tune the level of ergosterol (a molecule that is structurally similar to cholesterol) in their membranes. We find that reducing sterol levels in vacuole membranes causes them to phase separate, in contrast to previous assumptions.

## INTRODUCTION

Spatial organization of cellular components is essential for their biological function. One way that cells may achieve this organization is through spontaneous demixing of components into coexisting liquid phases. For example, subcellular condensates or droplets result from the demixing of proteins and/or RNA (reviewed in (1–5)). Similarly, model membranes phase separate to form 2-dimensional domains, independent of whether the membranes are composed of very few lipid types or are derived directly from cells (6, 7).

To date, yeast are the only unperturbed, living cell system in which reversible, micron-scale phase separation of a membrane has been observed. Specifically, domains disappear in individual yeast vacuoles above a characteristic temperature (*T*_mix_) and reappear below that temperature. Moreover, domains merge with one another on timescales characteristic of liquid membranes (8). This phase separation had been previously hypothesized (9, 10) and appears to regulate the cell’s metabolic response to nutrient limitation through the TORC1 (Target Of Rapamycin Complex 1) pathway that is a central regulator of protein synthesis, autophagy, docking of lipid droplets, and many other processes (10–14). Depletion of nutrients from the growth medium induces yeast to transition from exponential growth (called the logarithmic or “log” stage) to a stage in which cell division slows and the density of the cell culture plateaus (called the stationary stage, Fig. 1). When yeast enter the stationary stage, domains that segregate lipids and proteins appear on vacuole membranes, as observed by both freeze-fracture electron microscopy (15, 16) and fluorescence microscopy (10, 13). Immunogold labeling proves that domains seen by the two methods are the same (17).

**Figure 1:**
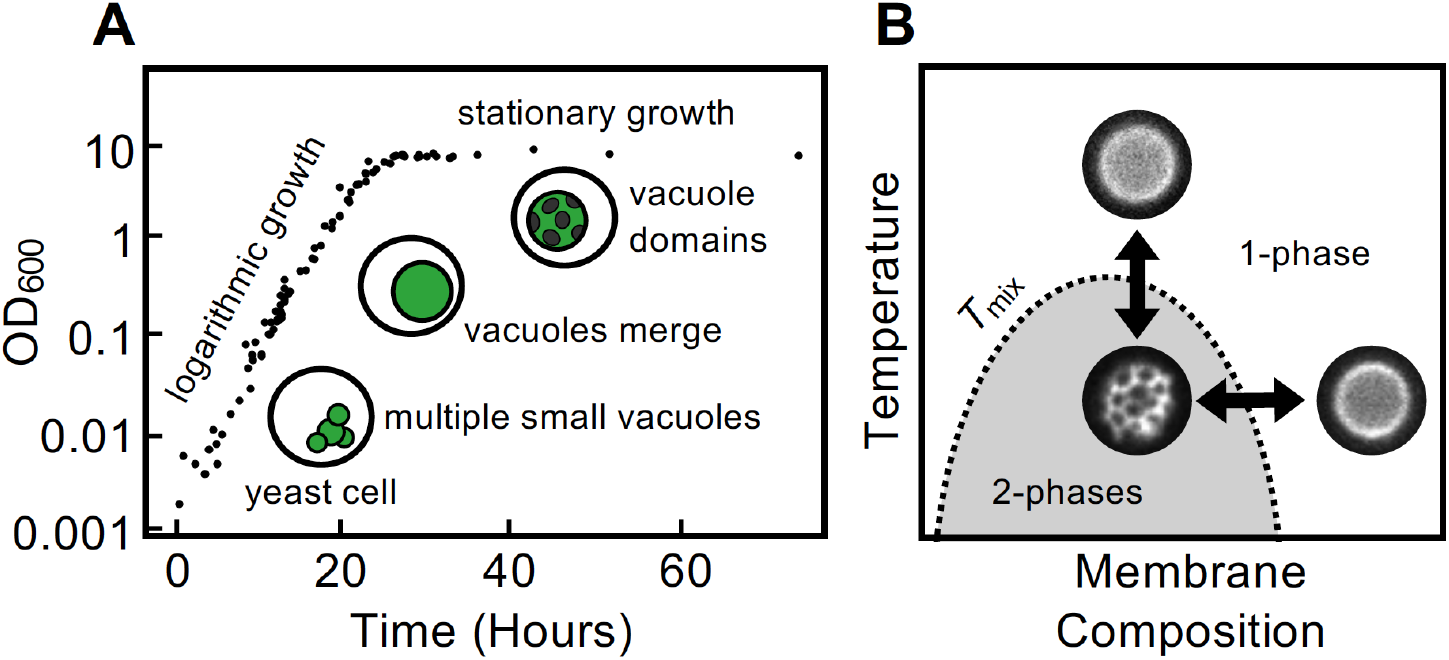
(A) Growth curve for *S. cerevisiae* (adapted from Rayermann et al. 2017), where OD_600_ is the optical density at λ = 600 nm plotted on a log scale. In the logarithmic stage of growth, yeast vacuole membranes appear uniform, in a single liquid phase. Late in the log stage, vacuoles fuse so that most cells contain only one large vacuole. In the stationary stage, the vacuole membrane demixes into two liquid phases with compositionally distinct domains. (B) The phase boundary between mixed (1-phase) and demixed (2-phase) states can be reversibly crossed by changes in temperature and/or membrane composition. *T*_mix_, the temperature at which membrane domains appear or disappear, changes with membrane composition. Micrographs show representative images of membranes with and without domains (and are not all from the same vacuole).

The clear connection between membrane domains and growth stage makes yeast an ideal system to investigate in order to answer significant open questions about phase separation. Here, we focus on two outstanding questions. First, do yeast cells physiologically adapt to maintain membrane phase separation under new physical conditions? Second, is phase separation induced or eliminated through manipulation of sterol levels?

It is reasonable to expect that yeast should actively remodel their vacuole membranes to achieve phase separation, because domains appear and disappear with growth stage and because yeast adjust to a wide range of environmental conditions (18, 19). More broadly, it is known that many cell types alter their lipid compositions and membrane behaviors in response to changes in cell cycle (20–25), activation (26, 27), disease (28–30), and environmental stress (31). Changes in growth temperature are particularly compelling to investigate because they are experimentally tractable and because they trigger active membrane remodeling in organisms ranging from bacteria to vertebrate animals (32–35).

If liquid-liquid membrane phase separation is a functionally relevant property, a clear prediction is that the membrane’s *T*_mix_ should be regulated by the cell in response to the ambient temperature. There is some precedent for this. Giant plasma membrane vesicles that are shed from zebrafish cells exhibit fluctuating domains indicative of a miscibility critical point, and the *T*_mix_ values of these cell-free membranes vary with the growth temperature of the cells that shed them (35). However, there are key differences between plasma membrane vesicles and yeast vacuoles, beyond the fact that one is cell-free and the other is in a living cell. To date, the *T*_mix_ of all plasma membrane vesicles has been found to be below the cells’ growth temperature (7). In these vesicles, an offset between *T*_mix_ and the growth temperature is hypothesized to set the size of submicron membrane fluctuations, which could in turn influence channel activities and cytoskeletal compartmentalization (36–38). In contrast, the *T*_mix_ in living yeast vacuoles is well above the growth temperature (8), such that vacuole membranes partition into micron-scale domains under normal conditions. In yeast, it is less clear why a specific offset would be worth regulating: any *T*_mix_ above the growth temperature should produce large domains in vacuole membranes. However, by regulating an offset, yeast might avoid “burning their bridges”, facilitating a return to a uniform membrane.

Of course, yeast cells cannot regulate domain formation by controlling their own temperature. Instead, they must alter the molecules in their membranes. Asking whether yeast tightly control their membrane’s *T*_mix_ is the same as asking whether they tightly control their lipid and protein compositions in the stationary stage (Fig. 1B). The detailed mechanisms by which cells adapt their membrane compositions to invoke phase separation in the stationary phase is unknown. In model membranes, coexisting liquid phases appear at intermediate sterol levels (rather than at low or high levels) (6, 39–43). In yeast cells, there are many indications that sterol levels are important. Experiments that alter genes encoding proteins involved in sterol synthesis or transport (11–13, 17), that perturb lipid droplets (which are rich in sterol esters) (13, 14), and that manipulate sterol metabolism (10, 44) all support the hypothesis that sterols regulate the formation and maintenance of membrane domains. However, it is largely unknown in which direction sterols are being shuttled—to or from the vacuole membrane.

In order to address some of these outstanding questions, we investigated whether living yeast cells adjust the *T*_mix_ of their vacuole membranes to scale with the growth temperature of the cells, and we test whether vacuole domains appear or disappear when ergosterol levels are changed in isolated vacuole membranes.

## RESULTS

In previous experiments (8), we investigated a relatively small number of individual vacuole membranes, both within and isolated from living cells. Here, we investigate large populations of cells. Three scenarios are possible. (1) If phase separation is physiologically irrelevant, then we would predict that *T*_mix_ is not stringently regulated: different cells within a population should exhibit *T*_mix_ values that range widely above and below the growth temperature. (2) If phase separation is functionally important, and the only constraint is that the vacuole membrane must phase separate in the stationary phase, then we might predict that membranes from different cells should have a wide range of *T*_mix_ values, as long as *T*_mix_ is usually higher than the growth temperature. (3) However, if phase separation is functionally important, and if the phase transition must be easily reversed when nutrient conditions change, then we would predict that *T*_mix_ is tightly regulated, and that the distribution of *T*_mix_ values should exhibit low cell-to-cell variation across the population.

### Yeast vacuoles exhibit a characteristic T_mix_

Vacuoles in living yeast cells were imaged as in Fig. 2A. The bright areas of the membranes are labeled with Vph1-GFP. Vph1 localizes exclusively to the vacuole membrane and is a transmembrane subunit of the V-ATPase pump. We chose Vph1 because it preferentially partitions to liquid disordered domains in phase-separated vacuoles (10, 17, 45). The protein’s function, which is to pump protons into the vacuole lumen to generate an electrochemical gradient across the membrane, is not affected by the probe (46). At temperatures below *T*_mix_, Vph1 is unevenly distributed in vacuole membranes (marked with arrowheads in Fig. 2A), whereas above *T*_mix_ the membranes appear uniform. To obtain the data in each experiment, multiple fields of view as in Fig. 2A were analyzed with a semi-automated, blinded procedure. For a population of cells, *T*_mix_ is the temperature at which the percent of vacuoles with domains is reduced by 50%. For the single experiment in Fig. 2B, each data point represents hundreds of vacuoles in living yeast cells grown at their optimal growth temperature of 30°C.

**Figure 2:**
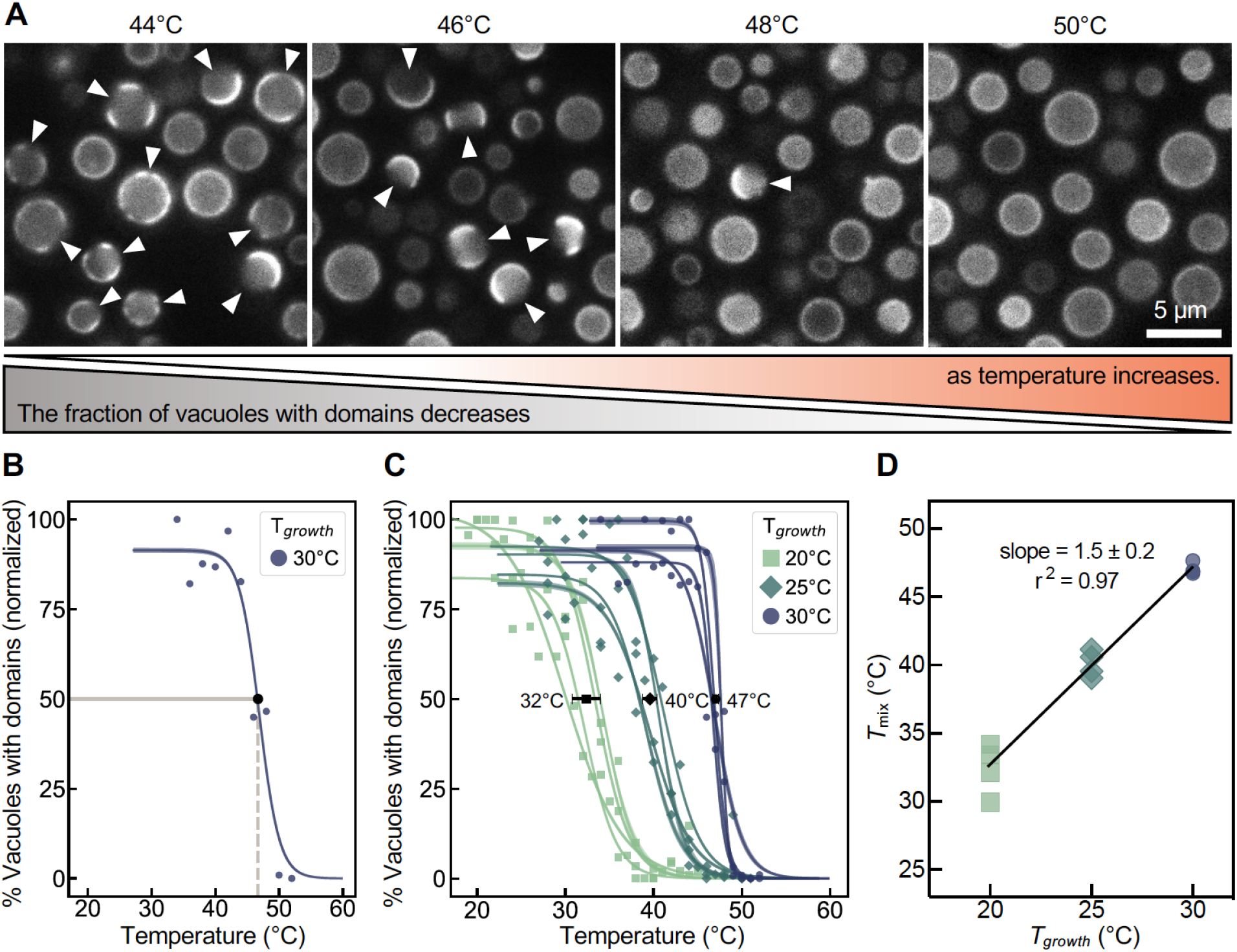
(A) Micrographs of vacuoles in living cells at 44°C, 46°C, 48°C, and 50°C, in four different fields of view. At low temperatures, most vacuole membranes are phase separated, whereas at high temperatures, most membranes appear uniform. Arrowheads point to membrane domains. (B) Vacuole membranes in living cells have a characteristic miscibility transition temperature, *T*_mix_. This plot represents a single experimental sweep with 2°C steps for cells grown at 30°C. The percent of vacuoles with visible domains (small circles; n = 235 to 466) was scored using a semi-automated blind procedure, normalized to a maximum of 100%, and fit to a sigmoidal curve (see Methods and Fig. S5). The large black circle lies at *T*_mix_ = 46.9°C, the temperature at which the percent of vacuoles with domains is reduced by 50%. The shaded region superimposed on the line indicates the 95% confidence interval for the fit. (C) Increases in growth temperature (20°C, 25°C, and 30°C) result in increases in *T*_mix_ (32.4 ± 1.6°C, 40.1 ± 0.8°C, and 47.0 ± 0.4°C, where error bars are standard deviations). Each curve represents a different experimental temperature sweep. (D) The relationship between *T*_mix_ and the growth temperature is linear. The error in the slope is the 95% confidence interval for the linear regression fit. Offsets (Δ*T*) between the three growth temperatures and *T*_mix_ are 12.4 ± 1.6°C, 15.1 ± 0.8°C, and 17.0 ± 0.4°C.

In turn, multiple experiments as in Fig. 2B were combined to generate Fig. 2C. We find that the broad population of vacuoles grown at 30°C has a characteristic *T*_mix_ (47.0°C) with a low variance (standard deviation ±0.4°C). Because *T*_mix_ is a function of the membrane’s composition, the sharp transition implies that there is relatively little cell-to-cell variation in vacuole membrane composition with respect to molecules that control phase separation, even among different yeast cultures. The reproducibility of this result rivals those of chemically defined artificial lipid systems like giant unilamellar vesicles (43, 47). In summary, yeast regulate their vacuole membrane composition to achieve a common *T*_mix_, which is set above the ambient growth temperature.

### Growth temperature controls T_mix_

Next, we tested whether cells actively tune *T*_mix_ of their vacuole membranes in response to changes in ambient growth temperature. We chose the new growth temperatures to be low (20°C and 25°C) rather than high to avoid subjecting cells to chronic thermal stress. Yeast grow robustly at all temperatures tested (Fig. S1 and Methods).

The results, shown in Fig. 2C and 2D, indicate that yeast maintain a linear relationship between *T*_mix_ in vacuole membranes and the growth temperature. An increase in the ambient temperature of ~15°C causes membrane components to mix into a single phase. Taken together, the low cell-to-cell variation of *T*_mix_ and the linear relationship between *T*_mix_ and the growth temperature strongly support the conclusion that vacuole membrane demixing is stringently regulated and, therefore, physiologically important.

### Membranes require long-term changes in temperature to remodel

Next, we investigated the timescales over which temperature changes regulate domain formation. Previous results indicate that the relevant window falls between 5 minutes and 3 hours. The shorter time scale derives from our previous report that domains do not reappear in vacuoles held above *T*_mix_ for 5 minutes (8). The longer time comes from the observation that domains eventually appear in log-stage vacuoles subjected to 3 hours of acute heat stress (10). We grew cells at low temperature (20°C) until they reached stationary stage and their vacuoles exhibited domains (Fig. 3). We then raised the temperature slightly above *T*_mix_ (35°C). We maintained the cells at this new temperature for 1 hour. Within this period, vacuole membranes did not return to a phase-separated state. Therefore, the time required for significant remodeling of the membranes is > 1 hour. We designed our experiment to avoid heat stress by choosing an initial low temperature (20°C), which resulted in a low *T*_mix_ (32.4 ± 1.6°C). At the conclusion of the experiment, cells were cooled back to 20°C, and domains reappeared on vacuoles that originally exhibited them (Fig. S2). We previously showed that miscibility transitions in vacuoles are reversible after acute changes in temperature on timescales of seconds (8). Here we show that the transition is also reversible on a long time scale, suggesting that *T*_mix_ is regulated in response to a time-averaged growth temperature, rather than *via* rapid cellular responses to acute temperature fluctuations.

**Figure 3:**
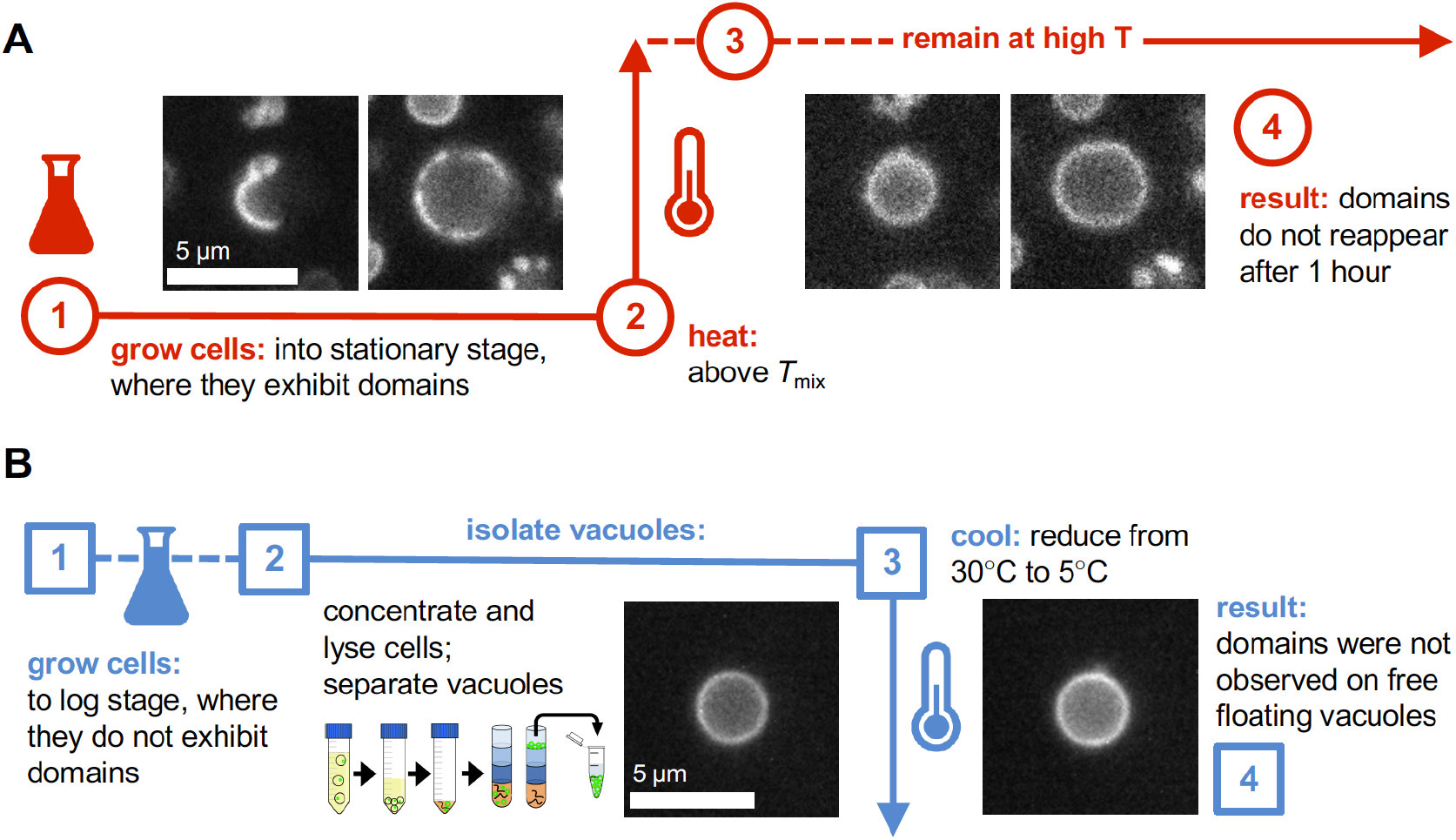
(A) Cells adapt *T*_mix_ of their membranes on timescales longer than 1 hour in response to temperature changes. Cells were grown at low temperature (~20°C) until they reached the stationary stage, where vacuole membranes exhibited domains. Temperature was then quickly raised to 35°C (slightly above *T*_mix_), and domains disappeared. Cells were maintained at this new temperature for 1 hour; domains did not reappear within this period. Images in Steps 1 and 4 show the same vacuoles; Fig. S2 shows a time lapse. (B) Membranes of free-floating vacuoles isolated from cells in the log stage do not phase separate, even at low temperature. Cells were grown at 30°C to the log stage (OD_600_ = 1). Vacuoles were isolated from these cells and imaged. When the vacuoles were cooled from 30°C to 5°C, domains did not appear in the membranes. Images in Steps 3 and 4 show two different vacuoles that are representative of the population.

### Logarithmic stage vacuoles do not phase separate, even at low lemperature

As yeast progress from the log stage to the stationary stage, their vacuole membranes demix into two phases. This transition must be driven by a change in the membrane composition, but what is the magnitude of that change? If log-stage yeast require only a small change in the composition of their vacuole membranes to phase separate, then only a small drop in temperature should have the same effect. In Fig. 3b, we find that even a large drop in temperature (from 30°C to 5°C) does not cause log-stage vacuole membranes to phase separate, implying that the membranes undergo a large change in composition from log to stationary phase. For this experiment, we used isolated vacuoles that were fused to ~4 μm diameter because phase separation can be difficult to identify in smaller (~1-2 μm) *in vivo* log-stage vacuoles and because isolation does not appear to perturb phase separation (8–10). We imaged vacuoles that were free-floating rather than resting on the coverslip because it is known that contact between membranes and surfaces can promote membrane phase separation (48–50). Indeed, contact between isolated vacuoles and glass surfaces occasionally caused domains to appear in very cold (5°C) membranes (Fig. S3).

### Depletion (not addition) of ergosterol causes domains in log-stage vacuole membranes

In model membranes, coexisting liquid phases form when the membranes contain moderate amounts of sterol. If the sterol content is low, solid (gel) domains appear; if the sterol content is high, the membrane is uniform (6, 39–43). Given that no domains appear in membranes of log-stage vacuoles, one reasonable hypothesis is that their sterol content is too high to support phase separation. In this scenario, a decrease in the sterol content of log-stage vacuole membranes would cause coexisting liquid phases to appear. In yeast, the predominant sterol is ergosterol; it constitutes roughly 7-15 mole % of lipids in mid- or late-log stage vacuolar membranes (51, 52). In Table S1, we show that depleting ergosterol in model membranes of giant unilamellar vesicles can indeed increase *T*_mix_, whereas supplementing ergosterol can decrease *T*_mix_.

We tested our hypothesis by decreasing or increasing the amount of ergosterol in isolated log-stage vacuoles fused to ~4 μm diameter. To directly tune the membrane’s ergosterol content, we used methyl-β-cyclodextrin (MβCD) as a molecular shuttle to remove and add ergosterol. We found that a decrease in membrane ergosterol caused domains to appear in ~40% of log-stage vacuoles, whereas an increase had no effect (Fig. 4). The result is robust across several ratios of MβCD:ergosterol, showing that both gentle and stringent methods of ergosterol depletion are effective. Contact between membranes and surfaces can promote membrane phase separation (48–50), and when vacuoles rested on glass, domains nucleated at that surface if ergosterol was depleted, but not if ergosterol was supplemented.

**Figure 4:**
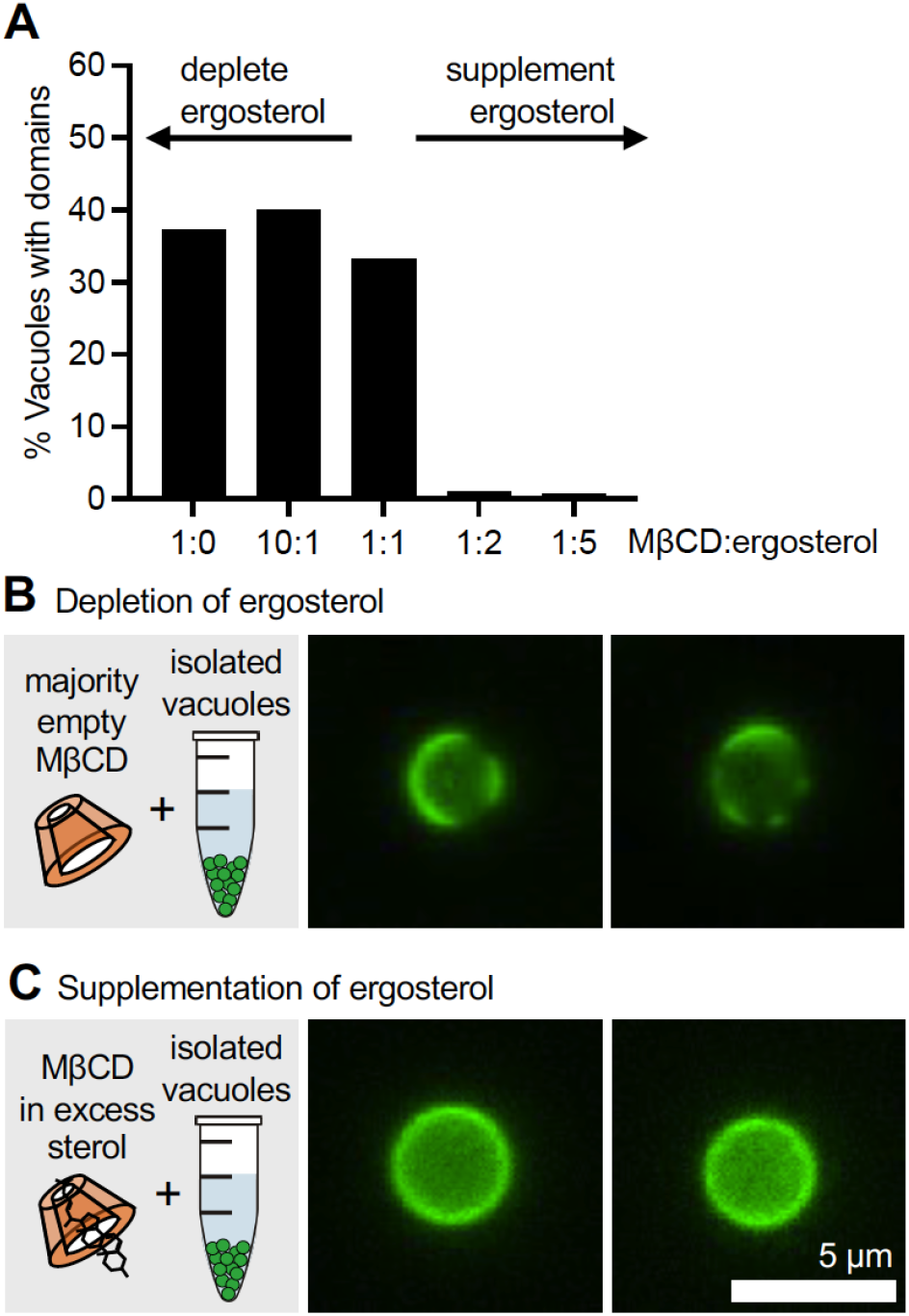
(A) In wild-type conditions, log-stage vacuoles do not exhibit domains (Fig. 1B). Depletion of ergosterol from vacuoles isolated from log-stage yeast triggers membrane phase separation, whereas supplementation of ergosterol has no effect. For each condition, 75 to 141 vacuoles were evaluated. (B) Micrographs show dark domains in isolated vacuoles in which ergosterol was depleted by introducing an excess of empty MβCD (as a 10:1 ratio of MβCD:ergosterol). Additional images are in Fig. S4. (C) Micrographs show a lack of micron-scale domains in isolated vacuoles in which ergosterol was supplemented by introducing loaded MβCD (as a 1:2 ratio of MβCD:ergosterol).

## DISCUSSION

### Phase separation in living cells

Cells expend energy to maintain molecular fluxes and concentration gradients far from equilibrium. A common question that arises about phase separation is how it can apply to living cells given that it is a classic equilibrium phenomenon. The apparent contradiction is reconciled if there is a clear separation of time scales. In vacuoles, the time required for the membrane to phase separate must be shorter than the time required for the membrane’s *T*_mix_ (or, equivalently, its composition) to change. We find that phase separation occurs within seconds of vacuole membranes crossing *T*_mix_ (8), whereas the value of *T*_mix_ remains constant (within measurement uncertainty) over timescales exceeding 1 hour (Fig 3A). A separation of time scales is also observed in subcellular condensates of proteins and/or RNA in living cells (1, 2, 4, 5). Timescales are also used in the classification of membrane domains. “Lipid rafts” are typically described as small (nm-scale), short-lived, dynamic platforms in plasma membranes (53–57), unlike the large (μm-scale), long-lived domains seen in vacuole membranes. Of course, phase separation could underlie many of the features attributed to rafts, such as segregating membrane components or influencing protein sorting and function (54, 58–60).

### The role of sterols in membrane phase separation

Here, we probed membrane composition by altering levels of only ergosterol, the main sterol in yeast. We changed sterol levels while keeping all other lipids constant by using MβCD and by isolating vacuoles from other types of membranes that could act as sources or sinks of sterols. We discovered that depletion of ergosterol causes log-stage vacuole membranes to phase separate.

Our experiments with vacuoles are consistent with results from artificial membranes. For many ratios of lipids in model ternary membranes, decreasing the sterol level is equivalent to traveling left along the horizontal arrow in Fig. 1b: domains nucleate and *T*_mix_ increases (e.g. (6, 41, 42)). A higher *T*_mix_ implies that a broader range of lipid compositions produces coexisting liquid phases (Fig. 1b). *In vitro* studies with cell-derived giant plasma membrane vesicles yielded similar results: decreasing sterol levels increases *T*_mix_ (35, 61). *In vitro* experiments with whole-cell lysates are more challenging to interpret. For example, Toulmay & Prinz reported that domains appear on log-stage vacuoles in lysates exposed to MβCD. Although their result agrees with ours, they surmised that MβCD was transferring sterols from other membranes in the lysate to vacuoles (10). Likewise, they found that MβCD caused domains to disappear from stationary-stage vacuoles (10), but it was not known if ergosterol was being transferred to or from the vacuoles.

Experiments that allow several lipid types to vary at once are similarly challenging to interpret. The list of lipids in yeast membranes is long and diverse, and changes in ergosterol levels could work in concert with (or against) changes in the rest of the lipidome to form membrane domains (62, 63). Most *in vivo* experiments imply that ergosterol levels in vacuoles increase as yeast enter the stationary stage. For example, Tsuji et al. (17) report that filipin staining of sterols in stationary-stage membranes is higher than in log-stage, with the caveat that staining was relative to the plasma membrane rather than absolute. Similarly, genetic impairment of sterol synthesis and transport (11–13, 17), perturbations to lipid droplets (which are rich in sterol esters (13, 14)), and the application of drugs to manipulate sterols (10, 44) can prevent the formation or maintenance of domains. However, in many of these processes, the net direction in which sterols are being transferred is unknown. Proof that phase separation in living vacuoles always follows an increase in ergosterol would be consistent with the observation that maintenance of phase separation requires lipophagy, which is expected to mobilize sterols from esters stored in lipid droplets (13, 14, 17, 64). Sterol transport may also occur at sites where vacuole makes contact with the ER or other organelles (44). We are currently conducting experiments with purified vacuole membranes to learn how their lipidome remodels during the cell cycle.

### Regulation of T_mix_

Our finding that the *T*_mix_ of yeast vacuole membranes is tightly regulated by the ambient growth temperature is consistent with the hypothesis that phase separation is functionally important. This leads to an important question: what downstream biological activities depend on the formation of phase-separated domains at the vacuole? Domains in vacuole membranes segregate protein complexes of the TORC1 nutrient sensing pathway, which regulates autophagy (10–12). In addition, vacuole domains appear to be necessary for the docking and consumption of lipid droplets, a process called lipophagy (13, 14, 17, 64). Both autophagy and lipophagy are essential for helping cells survive periods of nutritional limitation. The proposed role of membrane phase separation in the TOR signaling pathway may be especially significant, as the TORC1 pathway is conserved in yeast and humans.

Moreover, our findings imply that new mechanisms of regulating the composition of yeast vacuoles may await discovery. Signaling pathways that could control phase separation in vacuole membranes in response to changes in temperature are, at best, only partially characterized. In a candidate-based screen for mutants whose membranes cannot phase separate, Toulmay and Prinz identified genes in at least three signaling pathways: the Rim system, which responds to pH, the Fab1 (also called PikFYVE) phosphatidylinositol-3-phosphate 5-kinase, which responds to osmotic gradients and other stresses, and the Slt2 (Mpk1) MAP kinase, also involved in osmotic and membrane stress responses (10). At least two of these systems are also important for cellular responses to temperature. Mutant *fab1*Δ cells exhibit severe growth defects at elevated temperature, demonstrating that Fab1 activity is indispensable for cellular thermotolerance (65). Similarly, genes encoding proteins in the Slt2 pathway are mutated under selection for thermotolerance in experimental evolution studies, and this pathway becomes constitutively active during mild heat stress (66, 67). Moreover, we previously found that both phospholipid acyl chain composition and plasma membrane membrane fluidity are regulated through the Rho1–Protein Kinase C–Slt2 signaling pathway (68).

Phase separation in the yeast vacuole, first observed half a century ago, has re-emerged as the most robust and tractable model for understanding membrane phase separation in an intact cell system. Recent results from several groups show that essential processes for domain formation include lipophagy, sterol metabolism, and probably phospholipid metabolism. Our present results reveal an unexpectedly tight thermostatic regulation of the vacuole membrane’s biophysical properties. Going forward, major challenges include dissection of the mechanisms underlying this thermostat, and tests of the downstream functional consequences of membrane phase separation. Unresolved biophysical questions include the mechanisms that facilitate or impede the merging of many small domains into a few large domains in yeast vacuoles and the roles of transmembrane osmotic gradients and membrane tension in domain formation and structure.

## MATERIALS AND METHODS

### Yeast Cell Culture

*Saccharomyces cerevisiae* BY4742 (69) harboring a Vph1-GFP translational fusion was used in all experiments (*MATa his3Δ1 lys2Δ0 ura3Δ0 leu2Δ0 VPH1-GFP::HIS3*). Cultures were grown in a shaking incubator at 225 rpm using synthetic complete (SC) media containing 0.4% dextrose. For log stage experiments, cultures (1 L) were grown to an optical density @ 600 nm (OD_600_) of 1.0 (~17 hours at 30°C).

For stationary stage experiments, cultures (10 mL) were incubated at 20°C, 25°C or 30°C. Yeast were harvested after 72 hours of growth at similar cell densities (OD_600_ ≈ 6). Cultures at 20°C grow at a slower rate and therefore were started at a higher concentration to reach the same optical density (a proxy for growth stage) in an equivalent amount of time. We prioritized cell density because it has been shown to influence transition temperatures in cell-derived membranes (25, 26). Incubator temperature was monitored with a Fisherbrand TraceableLIVE Wi-Fi Datalogging Thermometer (0.25°C accuracy). This monitoring renders phase transition temperatures in this work more accurate than reported previously (8).

### Temperature Experiments

Cells were diluted for imaging in an isosmotic solution of conditioned media from the grown culture. To make conditioned media, 1 ml of culture was centrifuged at 3400×g. The supernatant was collected and centrifuged again at 3400×g rpm to remove remaining cells. To decrease refractive index mismatch (70), 200 μL of OptiPrep (60% OptiPrep Density Gradient Medium Sigma Cat # D1556) was added to 800 μL media and vortexed. Coverslips were coated with 3 μL of 1 mg/ml Concavilin-A (EPC Elastin Products Co no. C2131) in buffer [50 nM HEPES (pH 7.5, 20mM calcium acetate, 1 mM MnSO_4_]. Immediately prior to use, the coverslips were washed with MilliQ water and dried with air. Samples of 3 μL of cells were diluted into 3 μL of conditioned media containing 12% OptiPrep and placed onto the concanavalin-A coated coverslip. Cells were sandwiched with a top coverslip and allowed to adhere to the coated coverslip for 10 min before imaging.

Images were acquired on a Nikon TE2000 using a Blackfly 2.3 MP Mono USB3 Vision or Teledyne Photometrics Prime 95BSI camera. Fluorophores were excited with an X-Cite 110 LED light source filtered through an IR cut filter to prevent aberrant heating of the sample from the optics. A temperature cuff (Harvard Biosciences Inc.) on the oil immersion objective (100, 1.4 NA) was used to prevent the microscope from acting as a heat sink. Thermal grease (Omega OT-201) was used to keep the sample in thermal contact with the home-built, temperature-controlled stage (Omega Engineering). The sample temperature was measured using a thermistor probe (0.2°C accuracy, Advanced Thermoelectric). Temperature was adjusted stepwise in 2°C increments, starting at the growth temperature and increasing until domains were no longer observed. Temperature sweeps were done quickly, and only at high temperature for 3-5 min, to limit cellular thermal stress. Micrographs were taken at each 2°C interval and were used to quantify the fraction of vacuoles with domains at each temperature.

### Image Analysis

To preserve the fidelity of the data, image manipulation was limited to adjusting overall brightness and linear (γ=1) contrast enhancements, which was accomplished with ImageJ (http://imagej.nih.gov/ij/). To score membranes as being either uniform or phase separated, images were automatically cropped and displayed to the user using original Python code that has been made available by public license at github.com/leveillec/demixing-yeast-vacuole. The user was blind to the growth and sample temperatures for each image (Fig. S5). The percent of phase separated vacuoles at each temperature was fit using

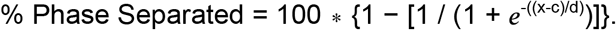

Here, *c* is *T*_mix_, the temperature at which the percent of vacuoles with domains is reduced by 50%, and *d* is the rate of the sigmoid decay.

At all temperatures, at least some vacuole membranes appear uniform, as reported by others (9, 10, 13). Data were normalized by setting the maximum percent of phase separated vacuoles to 100% for each experiment. The experiments systematically undercount vacuoles with domains because micrographs capture only one plane of the vacuole, which may not intersect with a domain. Our data also undercount vacuoles in which domains are too small to identify; excess area in the membrane can prevent domains from merging (71). During the course of an experiment, domains can coalesce (Fig. 2A), rendering them easier to identify and leading to an increase in the apparent percentage of vacuoles with domains as *T*_mix_ is approached from low temperatures.

### Vacuole Isolation Experiments

A 1 L culture was grown to OD_600_ = 1.0. Cells were sedimented in a swinging bucket rotor for (3200×g, 10 min, room temperature), resuspended in 0.1 M Tris (pH 9.4) and 10 mM dithiothreitol, and incubated for 10 min at 30°C. The cells were again sedimented in a swinging-bucket rotor (3200×g) for 5 min at room temperature and then resuspended in spheroplast buffer (600 mM sorbitol, 50 mM potassium phosphate pH 7.5, and 8% growth media). Lytic enzyme (Zymolyase 20T, Seikigaku; further purified in-house by cation exchange chromatography) was added and the cells were incubated for 25 min. The spheroplasted cells were sedimented in a swinging-bucket rotor (3200×g) for 5 min at 4°C. For hypoosmotic lysis, spheroplasts are resuspended in 15% ficoll buffer (10 mM Pipes-KOH pH 6.8, 200 mM sorbitol, and 15% w/v ficoll) and diethylaminoethyl-dextran was added to a final concentration of 0.005–0.01% w/v. Spheroplasts were incubated for 2 min on ice, then 3 min at 30°C. The resulting spheroplast lysates were added to a SW-41 ultracentrifuge tube and overlaid with a step gradient of 8%, 4% and 0% Ficoll in PS buffer (10 mM Pipes-KOH, pH 6.8, 200 mM sorbitol). Ultracentrifugation in an SW-41Ti rotor (Beckman) at 30,000×g for 90 min at 4°C resulted in pure vacuoles at the 4%-0% Ficoll interface.

Images of cell-free vacuoles were collected by diluting 5 μL of the isolated vacuole prep in 45 μL of low-melt 0.8% agarose prepared in PS buffer and mounted between a microscope slide and coverslip. The agarose immobilized vacuoles during imaging and prevented them from “splatting” onto the coverslip. Images were collected on an Olympus IX71 fluorescence microscope with an Andor IXON electron-multiplying charge-coupled device camera, using a 60 1.45 NA oil immersion objective and 1.5 tube lens.

### Cyclodextrin:Ergosterol complex

Ergosterol was dissolved in chloroform in a glass test tube and the solute chloroform was evaporated using a gentle stream of dry N_2_ to create a thin film. The film was then rehydrated in 2.5 mM methyl-β-cyclodextrin in aqueous solution. The tube was vortexed and then sonicated in a bath sonicator for 1-3 min. The solution was incubated in a 37°C water bath overnight and used the next day.

## ACKNOWLEDGMENTS

We thank Rachel Plemel, Tom Duan, and Scott Rayermann for technical advice. This research was supported by National Science Foundation grant MCB-1925731 to S.L.K., National Institutes of Health (NIH) grants R01 GM077349 and GM130644 to A.J.M., and NIH fellowship T32 GM008268 to C.E.C.

