## Supplementary Material for "Yeast cells actively tune their membranes to phase separate at temperatures that scale with growth temperatures"

**This PDF file includes:**

Supplementary Methods
Figures S1 to S6
Table S1
SI References

SUPPLEMENTARY METHODS

**Giant unilamellar vesicles:**

GUVs were prepared with 60 mol% DOPC, 20 mol% DPPC, 20 mol% ergosterol, and 0.8 mol% Texas Red DPPE and electroformed as previously described (1). Briefly, an aliquot of 0.25 mg of lipids dissolved in chloroform was spread evenly on a glass slide coated with indium-tin-oxide (ITO, Delta Technologies, Loveland CO). The slide was placed under vacuum for 30 min to evaporate the chloroform. A capacitor was created by separating two ITO-coated slides with two rectangular Teflon bars (0.3 mm thick). The gap between the bars was filled with ultrapure (18 MΩ-cm) water and all edges were sealed with vacuum grease. An AC voltage of 10 Hz and 1.5 V was applied to the capacitor for 1 hour at 60ºC. Vesicles were then extracted from the capacitor and diluted 5–10-fold in ultrapure water.

To image GUVs, several drops of vesicle stock solution were deposited between glass cover slips, and the edges were sealed with vacuum grease. This assembly was thermally coupled to a temperature-controlled stage on a Nikon Y-FL epifluorescence microscope via a layer of thermal grease. Images were captured using a 10x air objective.

**Synthetic complete media recipe:**

Per Liter of $H_2O$ -
1.4 g Amino Acid Mix (see below)
1.7 g Yeast Nitrogen Base (without amino acids and ammonium sulfate)
5 g of Ammonium sulfate
4 g of Glucose (0.4% glucose) or 20 g of Glucose (20% glucose)

**Amino acid powder mix:**

| | |
|---|---|
| 1 g Adenine | 4 g Tryptophan |
| 1 g Histidine | 4 g Leucine |
| 1 g Methionine | 4 g Isoleucine |
| 1 g Uracil | 5 g Glutamic Acid |
| 1 g Arginine | 5 g Aspartic Acid |
| 2.5 g Phenylalanine | 7.5 g Valine |
| 3 g Lysine | 10 g Threonine |
| 3 g Tyrosine | 20 g Serine |

SUPPLEMENTARY FIGURES

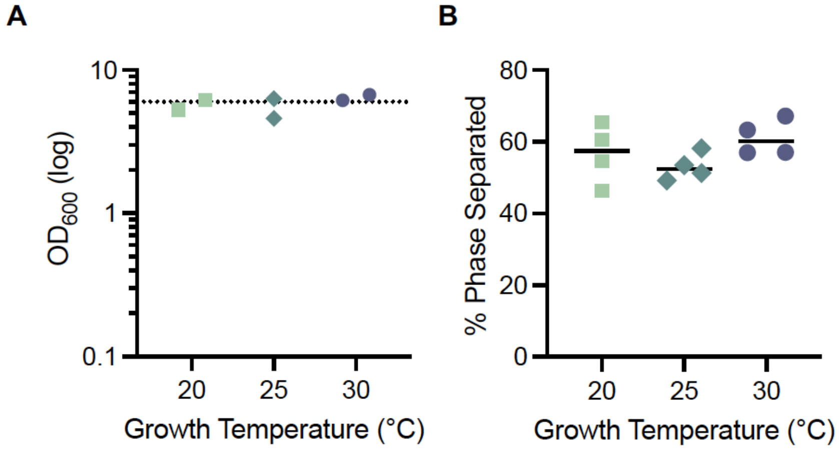


**Figure S1:** To control for the fact that growth temperature affects growth rate in yeast cells, we (A) grew cells to similar optical densities (OD) after 72 hours of growth, which produced similar percentages of vacuole membranes that were phase separated (shown in panel B). Yeast grown in 0.4% glucose reach the stationary stage after ~30 h at 25°C, and after ~12 h at 30°C. To achieve similar optical densities after 72 hours, starter cultures at 25°C and 30°C were diluted back to 0.001 OD. Cultures at 20°C were diluted back to 0.1 OD. If cultures at 20°C are diluted back to 0.001 OD, after 72 hours optical densities are only ~4, and only ~20% of the vacuoles have domains, indicating that the culture is not sufficiently into the stationary stage. (B) Less than 100% of vacuole membranes in the stationary stage exhibit domains, as reported previously (2–4). On average, cultures grown at 30°C exhibit slightly more phase separation in their membranes than cultures grown at cooler temperatures. For each population of cells in Fig. 2, the maximum percent of vacuole membranes with domains was normalized to 100%.

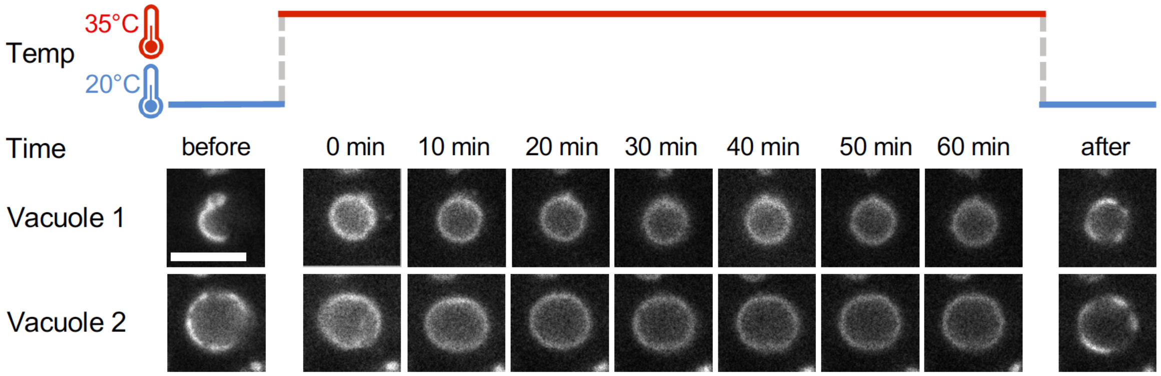


**Figure S2:** Timescales longer than 1 hour are required for yeast to significantly remodel their vacuole membranes. Yeast were grown at a low temperature (20°C) into the stationary stage, when their vacuoles exhibited domains with a $T_{mix}$ of 32.4 ± 1.6°C. The temperature was then raised to 35°C, just above $T_{mix}$. Cells were maintained at this new temperature for 1 hour. Within this time, vacuole membranes did not return to a phase-separated state. After 1 hour, the temperature was decreased to 20°C again, and domains were observed again. Images from two representative vacuoles are shown throughout the process. Scale bar = 5 μm.

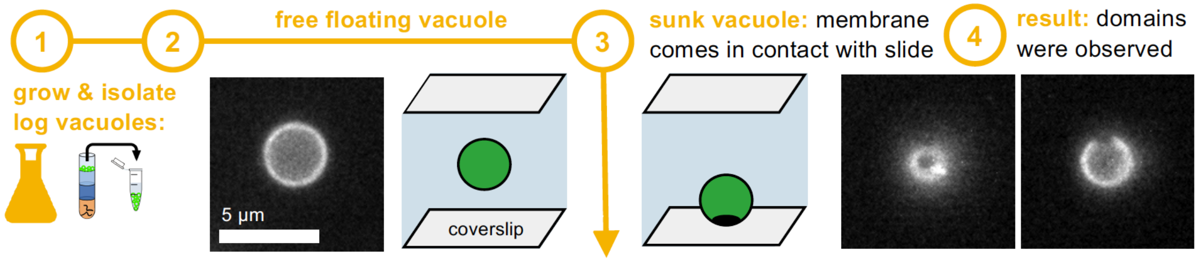


**Figure S3:** Vacuoles isolated from the logarithmic stage of growth do not exhibit domains. When cooled to 5°C, domains were not observed in the membranes of free-floating vacuoles. However, when the vacuoles came in contact with glass slides, domains were observed. This result is consistent with results in GUVs showing that domains can nucleate when the membrane is in contact with a surface and close to a phase transition (5–7).

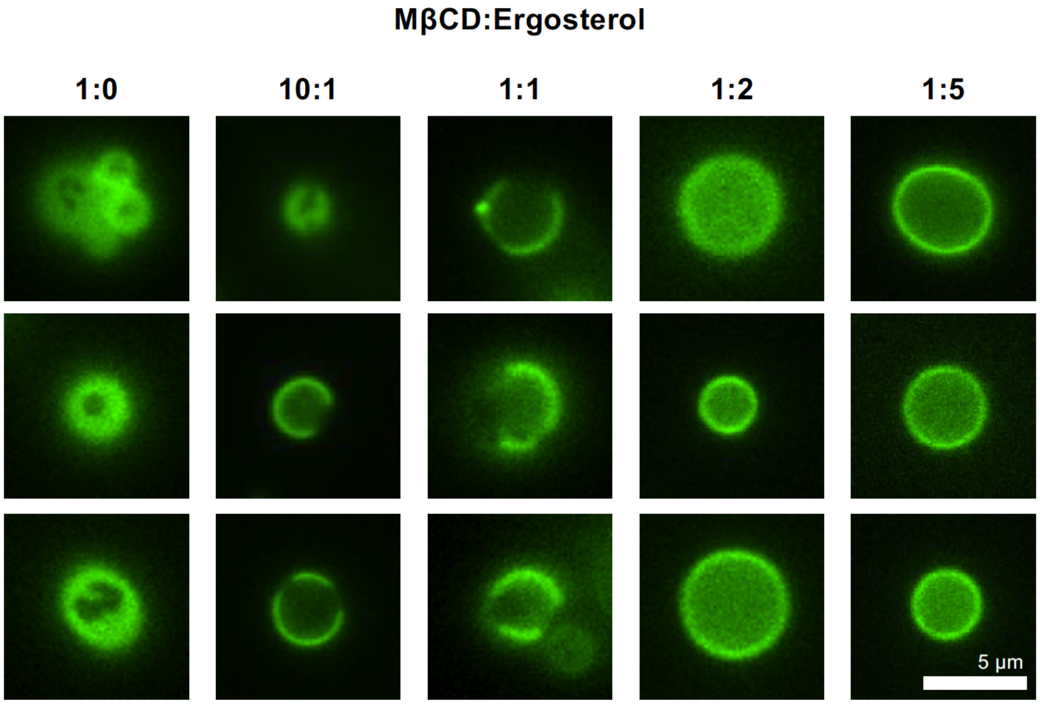


**Figure S4:** Domains appear in membranes of vacuoles isolated from yeast in the logarithmic stage of growth when ergosterol is depleted from the membranes. Depletion is achieved by introducing MβCD and ergosterol in ratios in which MβCD is in excess (1:0 and 10:1). When vacuole membranes are supplemented with ergosterol (by introducing MβCD and ergosterol in ratios in which ergosterol is in excess (1:2 and 1:5) domains do not appear – the membrane remains in a single phase. Introducing equimolar (1:1) MβCD and ergosterol has the same effect as depleting ergosterol from vacuoles.

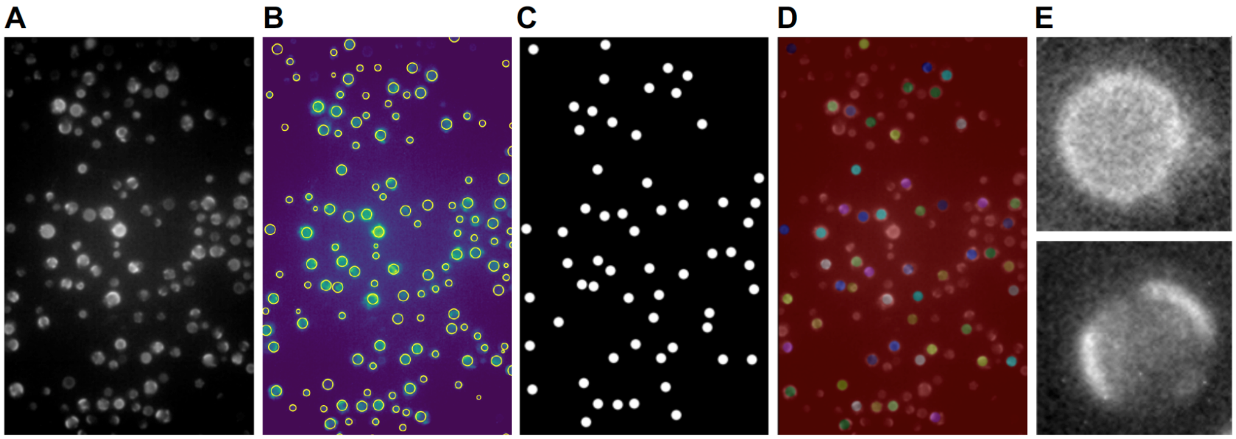


**Figure S5:** (A) Original images contained large fields of view. At each temperature, three fields of view were analyzed. (B) Individual vacuoles were identified as bright areas in the image using a blob detection "difference of Gaussians" function (www.scikit-image.org). (C) A mask was created using the coordinates and the radius from the blob detection. Vacuoles that were too small or along the edges of the image were discarded. (D) The vacuoles that remained were identified and individually cropped. (E) Each cropped vacuole was displayed and scored by the user as mixed (top) or demixed (bottom). The scorer was blind to the growth and sample temperatures for each cropped image. The scorer was also unable to see the whole field of view, so the temperature could not be easily deduced. Original Python scripts and the images above are available by public license at github.com/leveillec/demixing-yeast-vacuole.

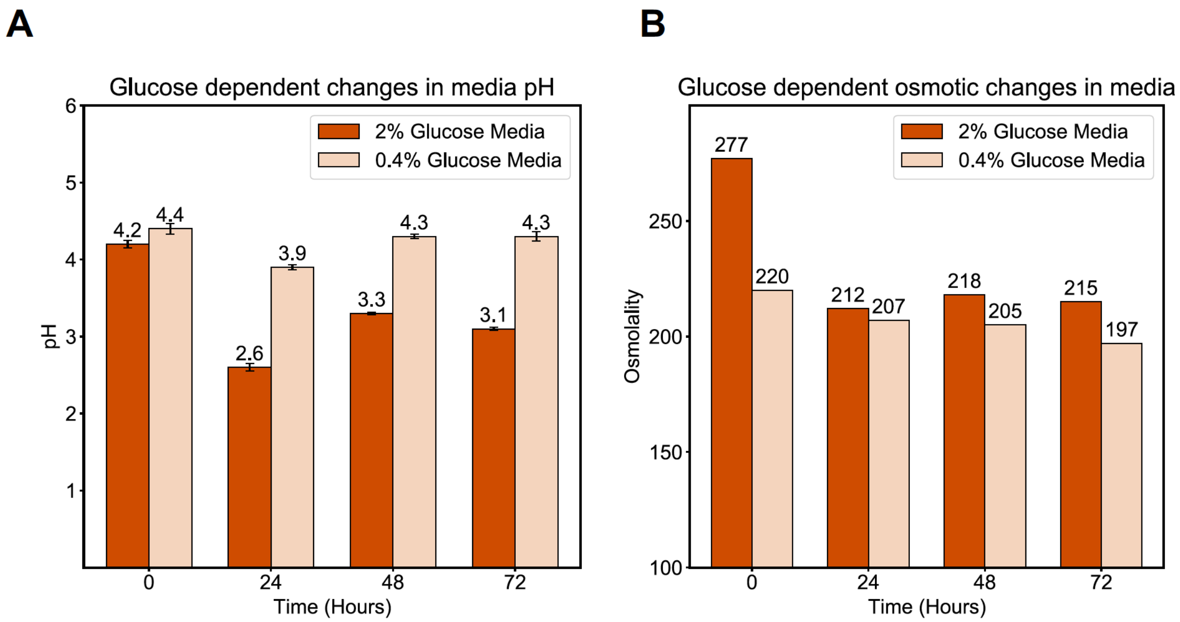


**Figure S6:** Media conditions are important to control because phase separation in vacuole membranes can occur in response to glucose depletion, acid stress, and osmotic shifts (2). (A) To minimize confounding effects from different stressors, we used SC media with 0.4% glucose, as opposed to 2% glucose because the pH of the media does not vary significantly through time when yeast are grown in 0.4% glucose. In contrast, when yeast are in media with 2% glucose, a large decrease in pH is observed as the cell metabolizes glucose into toxic species, including acetic acid (8). Error bars show the standard deviation of 3 measurements of the same sample. (B) Another reason that SC media with 0.4% glucose is preferable is that it does not result in a large osmotic shift during growth, whereas 2% glucose does. This experiment was performed once.

SUPPLEMENTARY TABLE

| Composition | Measured $T_{mix}$ | Change | Membrane ergosterol |
|---|---|---|---|
| GUV: 60/20/20 - DOPC/DPPC/Erg | 26.2 ± 0.3°C | – | Initial conditions |
| 10:1 MβCD:ergosterol added to GUVs | 29.4 ± 0.4°C | Increase in $T_{mix}$ | Membrane ergosterol depleted |
| 1:1 MβCD:ergosterol added to GUVs | 26.2 ± 0.2°C | No change in $T_{mix}$ | Ergosterol was added and removed at the same rate |
| 1:2 MβCD:ergosterol added to GUVs | 21.8 ± 0.2°C | Decrease in $T_{mix}$ | Membrane ergosterol supplemented |

**Table S1:** Depleting ergosterol from giant unilamellar vesicles (GUVs) increases $T_{mix}$; supplementing ergosterol decreases $T_{mix}$. GUVs were made of 60 mole % dioleoyl- phosphatidylcholine (DOPC), 20% dipalmitoylphosphatidylcholine (DPPC), and 20% ergosterol. Depletion was achieved by introducing MβCD and ergosterol at a ratio in which MβCD is in excess (10:1). This level of depletion of ergosterol increased $T_{mix}$ by ~3°C. Supplementation was achieved by introducing MβCD and ergosterol at a ratio in which ergosterol is in excess (1:2). This level of supplementation decreased $T_{mix}$ by ~4°C.
